# Y-RNA and tRNA Cleavage by RNase L Mediates Terminal dsRNA Response

**DOI:** 10.1101/087106

**Authors:** Jesse Donovan, Sneha Rath, David Kolet-Mandrikov, Alexei Korennykh

**Affiliations:** Department of Molecular Biology, Princeton University Princeton, NJ 08544, USA

## Abstract

Double-stranded RNA (dsRNA) is a danger signal that triggers endonucleolytic degradation of RNA inside infected and stressed mammalian cells. This mechanism inhibits growth and ultimately removes problematic cells via apoptosis. To elucidate the molecular functions of this program and understand the connection between RNA cleavage and programmed cell death, we visualized dsRNA-induced degradation of human small RNAs using RtcB ligase-assisted RNA sequencing (RtcB RNA-seq). RtcB RNA-seq revealed strong cleavage of select transfer RNAs (tRNAs) and autoantigenic Y-RNAs, and identified the innate immune receptor RNase L as the responsible endoribonuclease. RNase L cleaves the non-coding RNA (ncRNA) targets site-specifically, releasing abundant ncRNA fragments, and downregulating full-length tRNAs and Y-RNAs. The depletion of a single Y-RNA, RNY1, appears particularly important and the loss of this Y-RNA is sufficient to initiate apoptosis. Site-specific cleavage of small ncRNA by RNase L thus emerges as an important terminal step in dsRNA surveillance.

## Introduction

Cytosolic dsRNA is a major immunogen recognized by the mammalian innate immune system. The dsRNA molecules are not abundant in healthy cells because double-stranded regions are actively disrupted by cleavage (White et al., 2014) and editing (George et al., 2016; Liddicoat et al., 2015) mechanisms. Overload of these mechanisms by stress (Leonova et al., 2013) or viral infections (Kawai et al., 2005) results in dsRNA buildup, activating the type-I interferon (IFN) response. The IFNs upregulate multiple protective genes, and the combined action of these genes and dsRNA synergistically triggers intracellular RNA cleavage by the innate immune endoribonuclease RNase L (Donovan et al., 2013; Silverman et al., 1988). The action of RNase L promotes apoptosis to eliminate cells with excessive dsRNA load, whereas the absence of RNase L renders cells apoptosis-resistant (Castelli et al., 1998; Zhou et al., 1997). The precise mechanism of the suicidal dsRNA-RNase L signaling is not understood.

RNase L is a regulated ~ 83.5 kDa endoribonuclease built around a kinase-homology domain (Han et al., 2014; Huang et al., 2014). RNase L is activated by the binding of a dsRNA-induced mammalian second messenger 2-5A (Dong et al., 1994; Han et al., 2012) and the RNase L**•**2-5A complex cleaves single-stranded RNA (ssRNA), but not dsRNA. RNase L therefore does not serve for dsRNA removal. Efforts to understand the functions of RNase L in dsRNA surveillance found that some of the antiviral activity may arise from viral ssRNA degradation (Cooper et al., 2014; Han et al., 2004; Iordanov et al., 2000). However, RNase L inhibits cell growth (Al-Haj et al., 2012) and initiates programmed cell death (Zhou et al., 1997) independently from infections, indicating that cleavage of mammalian RNA is the apoptosis driver. RNase L can directly cleave 18S/28S ribosomal RNA (rRNA) to inhibit protein synthesis (Cooper et al., 2014; Hovanessian et al., 1979), degrade groups of messenger RNAs (mRNAs) to inhibit cell adhesion and wound healing (Rath et al., 2015), and generate small signaling RNA fragments shorter than ~ 200 nt to amplify the IFN response (Malathi et al., 2007) and inflammation (Chakrabarti et al., 2015). It remains unknown whether RNase L activates programmed cell death via a general stress caused by processing of multiple RNAs or via a single, perhaps still unidentified apoptotic RNA switch. Using a custom RNA-seq library that captures fragments derived from cellular RNAs, we identify surprising RNA targets of the dsRNA response, including a ncRNA that has properties of such an apoptotic switch.

## Results

### dsRNA Triggers Biogenesis of Specific tRNA and Y-RNA Fragments with 2’,3’-Cyclic Phosphate Termini

To identify the RNA fragments released by the action of dsRNA and RNase L, we focused our analysis on cellular RNAs bearing a 2’,3’-cyclic phosphate. Mammalian RNAs acquire this modification during endonucleolytic cleavage by RNase L and other metal ion-independent ribonucleases (Cooper et al., 2014). The RNA fragments were detected using RNA sequencing (RNA-seq) with RtcB ligase, an enzyme that joins RNAs with a 2’,3’-cyclic phosphate to RNAs with a 5’-hydroxyl (Tanaka et al., 2011). RtcB RNA-seq allows identification of the cleaved RNA and mapping of the cleavage site with single-nucleotide accuracy (Fig. 1A). This approach is not biased toward RNase L and will detect RNA cleavage by any metal ion-independent endoribonuclease activated during dsRNA response. Here, we examine small RNAs (≤ 200 nt), the only RNA class that was intractable during the recent mapping of the RNase L pathway (Rath et al., 2015). The isolation of small RNAs provides an essential advantage to our analysis as it eliminates the background from the ribosomal RNA (rRNA) (Cooper et al., 2014).

**Figure 1.**
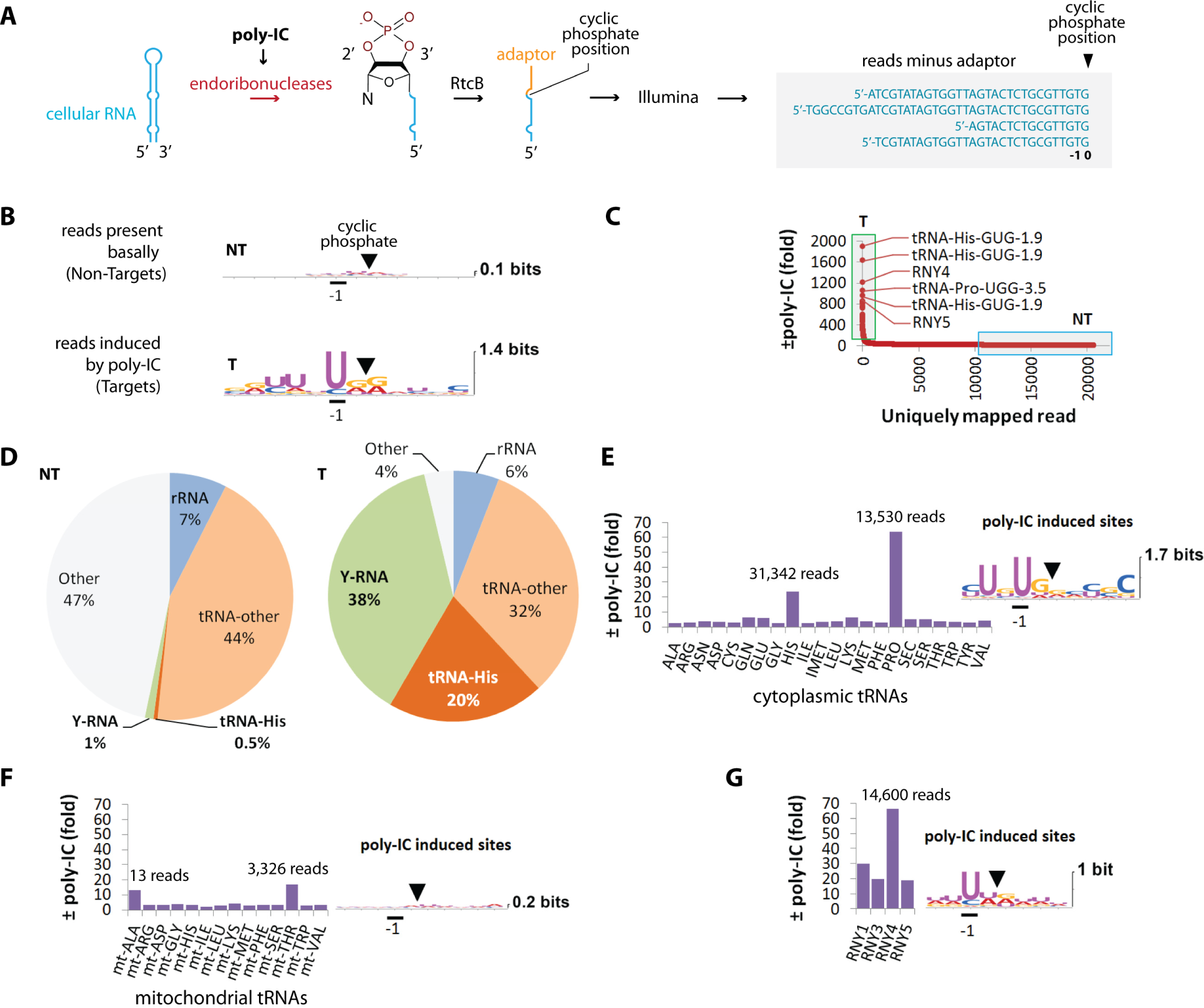
RtcB RNA-seq analysis of RNA cleavage in poly-IC-stimulated HeLa cells. (A) RtcB RNA-seq outline. (B) Nucleotide enrichments in RNA fragments from naive HeLa cells (non-targets, NT) and RNA fragments upregulated by ≥ 100-fold by poly-IC treatment for 24 hours (targets, T). (C) Distribution of uniquely mapping reads according to fold-upregulation. (D) Composition of non-targets and targets by RNA types. The group “other” contains fragments of small nucleolar RNAs, mRNAs, micro-RNAs and other small non-coding RNAs (Dataset S1). (E) Increase in the sum of fragments from each cytosolic tRNA upon poly-IC treatment. The cytosolic tRNAs have a strong UN^N consensus at cleavage sites. (F) Upregulation of mitochondrial tRNA fragments. Only tRNA-Thr cleavage is increased robustly and the UN^N consensus is absent. (G) Upregulation of the sum of Y-RFs by poly-IC.

RtcB RNA-seq in naive HeLa cells identified millions of basal reads with 2’,3’-cyclic phosphate and an average length of 9.4 nts (Fig. S1A). Transfection of a synthetic dsRNA (poly-inosine/poly-cytidine, poly-IC) released an abundance of new reads in the range of 10-70 nt, increasing the average read length to 18.3 nts (Fig. S1B). Reads that mapped unambiguously to the human transcriptome and had coverage depth of ≥ 5 reads (Dataset S1) were selected for all further analyses. Examination of the basal cleavage sites found no sequence enrichments (Fig. 1B). In contrast, poly-IC-induced cleavage sites had a distinct consensus, UN^N (^ marks the cleavage site), where N = A, G, U or C. Therefore, poly-IC activates a sequence-specific endoribonuclease, which requires a U at position −1. This requirement matches only RNase L (Han et al., 2014), indicating that other endoribonucleases do not contribute appreciably to the poly-IC-induced fragment production.

The most highly upregulated reads map to tRNAs and Y-RNAs (Fig. 1C). Fragments of tRNA-His and Y-RNAs account for more than 50% of the induced reads, whereas none of them are abundant basally (Fig. 1D). Fragments of tRNA-His and tRNA-Pro prevail among the cytosolic tRNAs (Fig. 1E), and fragments of tRNA-His accumulate to the highest absolute level compared to any tRNA. The tRNA cleavage consensus UN^N matches the specificity of RNase L. Notably, mitochondrial tRNAs lacked the UN^N consensus and neither mt-tRNA-His nor mt-tRNA-Pro was cleaved (Fig. 1F). The sequence-specific cleavage of cytosolic but not mitochondrial tRNAs argues against a possible mitochondrial action of RNase L (Le Roy et al., 2007).

All four human Y-RNAs, RNY1, RNY3, RNY4 and RNY5, are cleaved and contribute Y-RNA fragments (Y-RFs; Fig. 1G). The cleavage sites have a strong UN^N consensus, indicating the involvement of a single ribonuclease, presumably RNase L. These results reveal the molecular explanation for the recent finding of poly-IC-induced 22-36 nt fragments of RNY3 and RNY5 (Nicolas et al., 2012). The earlier study could not pinpoint the biogenesis mechanism, but concluded that the cleavage is Dicer-independent. Our data suggest that the cleavage is performed by RNase L and impacts not only RNY3/RNY5, but all Y-RNAs.

### Single-Nucleotide Resolution Profiling of tRNA and Y-RNA Cleavage

RtcB RNA-seq provides quantitative information about multiple RNA sites and allows visualization of detailed RNA cleavage profiles. This analysis shows that tRNAs and Y-RNAs are cleaved strongly at specific locations (Fig. 2). The cleavage of tRNA-His and tRNA-Pro occurs in the anticodon stem-loop (ASL) and results in tRNA halves. The formation of tRNA halves has been observed previously under cellular stress conditions and described as a mechanism important for sex hormone signaling (Honda et al., 2015), prostate and breast cancer proliferation (Dhahbi et al., 2014), and control of retroelements (Sharma et al., 2016). The primary route of tRNA half generation described to date involves cleavage of the ASL by the endoribonuclease angiogenin (Honda et al., 2015; Ivanov et al., 2014). Whereas angiogenin cleaves predominantly tRNA-Ala and tRNA-Cys between nucleotides 3/4 in the anticodon loop, poly-IC activates tRNA cleavage between nucleotides 4/5 (Pro) and 5/6 (His), releasing tRNA halves unique to dsRNA response.

**Figure 2.**
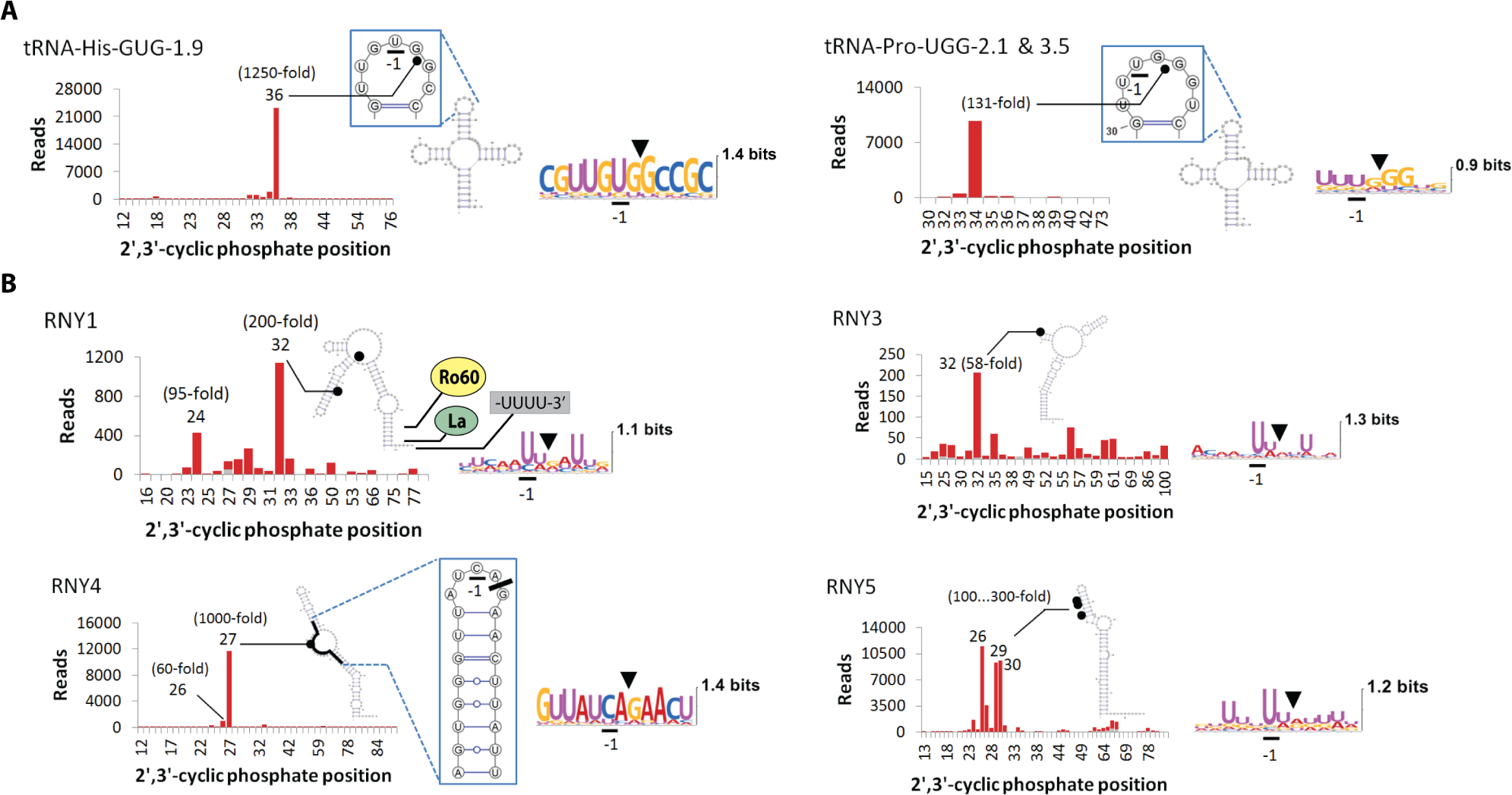
Single-nucleotide resolution profiles of tRNA and Y-RNA cleavage in poly-IC response. Stacked bar charts for RNA fragments detected by RtcB RNA-seq. Only positions with experimentally mapped cleavage sites are shown in the X-axis. Grey bars show read counts in na?ve cells. Red bars stacked on top of grey bars show read counts after poly-IC treatment for 24 hours. (A) Site-specific cleavage of tRNA-His and tRNA-Pro in the ASL. (B) Cleavage of Y-RNAs occurs in the upper region, which is not involved in the binding of Ro60 or La autoantigen proteins. RNY1, RNY3 and RNY5 are cleaved at UN^N sites; RNY4 is cleaved predominantly at a single site CA^G. The CA^G site is located in a single-stranded region, which may also form a local stem-loop structure. The most abundant Y-RFs arise from RNY5, followed by RNY4, RNY1 and RNY3.

The cleavage of Y-RNAs occurs at one or several closely located nucleotides and targets a narrow region demarcated by the nucleotides 24-32 (Fig. 2B). The four human Y-RNAs are evolutionarily related and contain ~ 100 nucleotides. They have an imperfect ~ 20 bp stem at the base and an oligo-U tail at the 3’-end (Kowalski and Krude, 2015; van Gelder et al., 1994) (Fig. 2B). The basal region of the Y-RNAs is structurally and biochemically characterized and serves as the binding site for at least two proteins, Ro60 and La (Alfano et al., 2004; Hung et al., 2015; Stein et al., 2005). The poly-IC induced cleavage sites identified here are located in the least understood, upper region of the Y-RNAs. This region binds, perhaps dynamically, different proteins (Kowalski and Krude, 2015).

Biomedically, Y-RNAs are the components of the autoantigenic complex that triggers autoimmunity in systemic lupus erythematosus (SLE) and Sjögren’s syndrome (Kowalski and Krude, 2015). Physiologically, Y-RNAs are essential for DNA replication, acting via an unknown mechanism that involves the upper stem (Gardiner et al., 2009). In addition, 22-36 nt Y-RFs are detected in the serum of cancer patients, in some normal cells and organs, and in apoptotic cells as complexes with Ro60 (Dhahbi et al., 2014; Kowalski and Krude, 2015; Rutjes et al., 1999). Some of these Y-RFs may arise from the action of RNase L because their size range overlaps with the size of the Y-RFs we observed here.

### RNY1/RNY5 Cleavage Requires Remodeling, RNY4 Defies the UN^N Consensus

The stereochemical mechanism of RNA cleavage via a 2’,3’-cyclic phosphate requires that the cleaved nucleotides are single-stranded (Korennykh et al., 2011). The identity nucleotide U in the UN^N pattern has to be single-stranded as well for binding to the specificity pocket in RNase L (Han et al., 2014). Therefore, RtcB RNA-seq reads arising from RNase L result from RNA substrates with single-stranded UN^N sites.

During the dsRNA response, RNY1 is cleaved predominantly at positions 24 (UA^U) and 32 (UU^G) (Fig. 2B). The first two nucleotides at position 24 are single-stranded and the third forms a weak base pair, which should permit cleavage via a 2’,3’-cyclic phosphate. In contrast, all nucleotides at site 32 are within a predicted RNA duplex and cannot be cleaved without a substantial structural remodeling in RNY1. In RNY3, the main cleavage site at nucleotide 32 (UA^A) resides in a stem-loop structure and is directly compatible with 2’,3’-cyclic phosphate formation (Fig. 2B). RNY4 is cleaved strongly and site-specifically. Global folding of RNY4 predicts that the single strong cleavage site is single-stranded. Local folding predicts a stable stem-loop structure (Fig. 2B), but still places the cleavage site within a single-stranded region compatible with 2’,3’-cyclic phosphate formation. Nonetheless, RNY4 is an outlier across the entire RtcB RNA-seq dataset. The cleaved sequence at position 27 (CA^G) violates the UN^N consensus, raising the question of whether the cleavage results from RNase L or another endoribonuclease. In RNY5, the cleavage sites 26 (UU^G), 29 (U^UA) and 30 (UA^A) are double-stranded and cannot result in a 2’,3’-cyclic phosphate without disrupting the RNY5 fold (Fig. 2B). Therefore, the proposed structure of RNY5 (van Gelder et al., 1994) must be remodeled at some point during the poly-IC response to release the target nucleotides from base pairing.

### The dsRNA-Induced tRNA Halves and Y-RFs Result Directly from RNase L

The UN^N RNA cleavage consensus observed during the poly-IC treatment implies that the responsible enzyme is RNase L. However, this implication is challenged by the paradoxical finding that RNY4 is cleaved at a CA^G site 27, whereas the flanking UN^N sites remain intact (Fig. 2B). To resolve this paradox, we used RtcB RNA-seq to profile RNA fragments in cells overexpressing RNase L. Overexpression of wild-type (WT), but not the inactive RNase L mutant H672N (Rath et al., 2015) induced tRNA and Y-RNA cleavage at sites with a strong UN^N consensus (Fig. 3A, B; Dataset S1). The cleavage of tRNA-His (129-fold) and tRNA-Pro (23-fold) occurred at the same ASL sites as in the poly-IC samples (Fig. S2A). The fold-induction of the tRNA fragments was smaller than using poly-IC treatment because overexpression activates RNase L mildly (Rath et al., 2015). RNY1 was not cleaved detectably due to the mild RNase L activation or a requirement for dsRNA for optimal RNY1 sensitivity. However, RNY3, RNY4 and RNY5 were cleaved at the same specific sites as in the poly-IC samples (Fig. S2B). These observations strongly suggest that tRNAs and Y-RNAs are cleaved by RNase L, including the non-canonical cleavage of RNY4.

**Figure 3.**
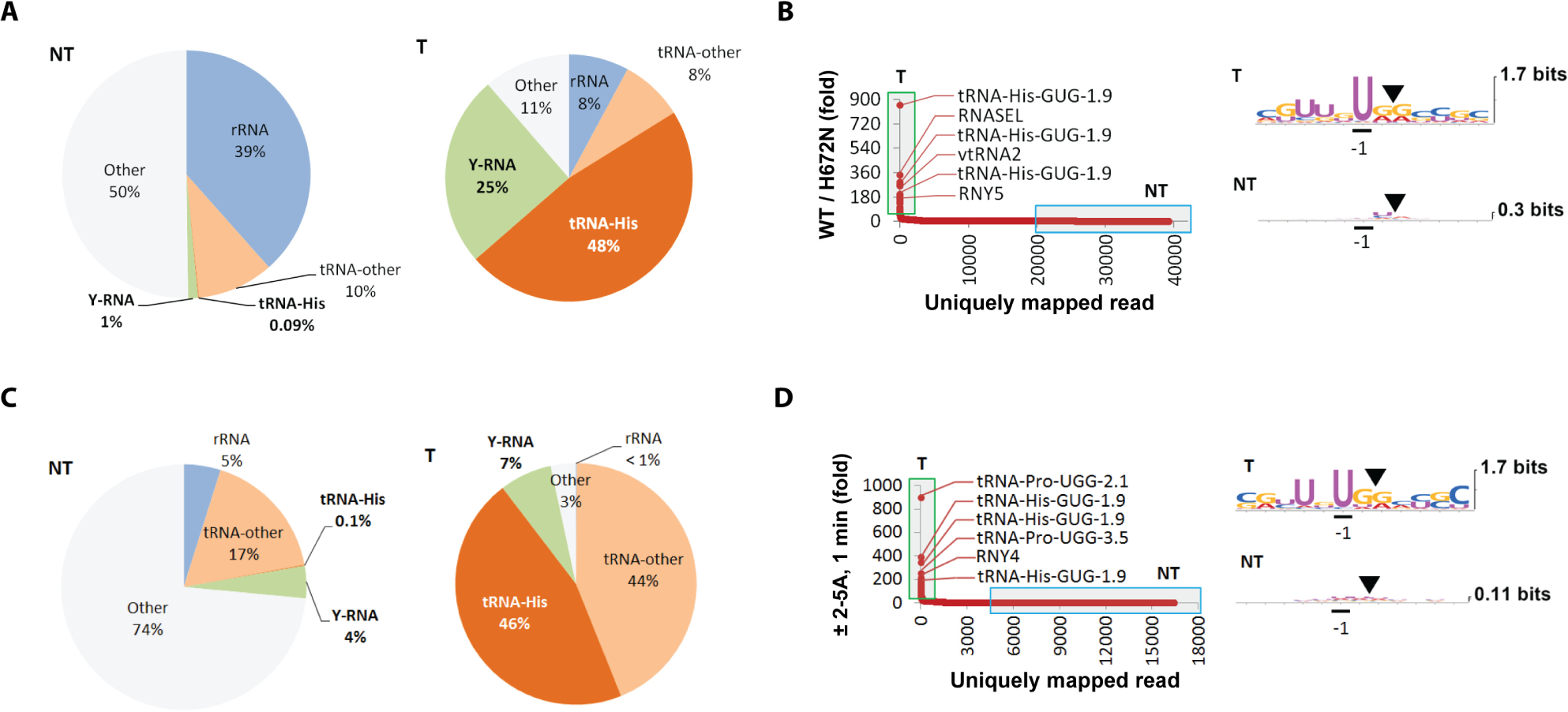
RtcB RNA-seq analysis of RNA cleavage in HeLa and T47D cells. (A) Composition of RtcB RNA-seq fragments upon overexpression of WT RNase L for 24 hours compared to H672N catalytically inactive mutant. The RNase L-induced change parallels the poly-IC response in figure 1D. (B) Distribution of uniquely mapped reads according to fold-induction. The cleavage sites produced by the action of RNase L have the UN^N consensus. (C) Composition of basal and 2-5A-induced RNA fragments in T47D cells treated with digitonin-2-5A. Fragments of tRNA-His and Y-RNAs account for more that 50% of the upregulated reads. (D) Y-RNA and tRNA fragments are upregulated by greater than 100-fold after one minute of endogenous RNase L activation. The cleavage sites show a strong UN^N consensus.

To more fully exclude secondary mechanisms, such as cleavage of RNY4 by an unknown, CA^G-specific endoribonuclease activated downstream of RNase L, we extended our RtcB RNA-seq analysis to semi-permeabilized T47D cells (Rath et al., 2015). This method employs 2-5A to activate endogenous RNase L very rapidly and allows assay completion within 1-3 minutes. The semi-permeabilization experiments revealed predominantly fragments of tRNAs and Y-RNAs, which were cleaved at the expected UN^N sites (Fig. 3C,D; Dataset S1). Analysis of tRNA cleavage revealed the same preference for tRNA-His and tRNA-Pro, as observed during poly-IC treatment (Fig. S3A; 1E). Both tRNAs were cleaved specifically at the correct ASL positions (Fig. S3B). The cleavage sites in Y-RNAs also exhibited the UN^N consensus, with the expected exception for RNY4, which was cleaved selectively at the CA^G position (Fig. S3C). The cleavage patterns of RNY3 and RNY4 resembled the patterns observed with poly-IC and RNase L overexpression, whereas the cleavage of RNY1 and RNY5 exhibited several differences. RNY1 was cleaved at the first poly-IC-induced site, 24, but not at the second site, 32 (Fig. S3C vs 2B), and exhibited a new cleavage site 66 (UU^G) located within a predicted stem-loop. RNY5 was not cleaved and exhibited only basally present fragments derived from RNase L-independent cleavage between positions 60-90 (Fig. S3C).

Strikingly, the altered cleavage patterns of RNY1 and RNY5 now agree better with the predicted secondary structures. The missing cleavage sites, 32 in RNY1 and 26, 29, 30 in RNY5, reside in RNase L-resistant RNA duplex regions. The conformational changes that must disrupt these duplex regions in RNY1 and RNY5 occur during poly-IC incubation or RNase L overexpression, but not during rapid 2-5A treatment. Apparently, the rearrangement in RNY1 and RNY5 involves a relatively slow step, which may include reorganization of cellular protein/Y-RNA complexes.

### Divergent Cellular Mechanisms are Essential for Y-RNA and tRNA Recognition by RNase L

Recognition of the UN^N sequence is insufficient to explain the preference of RNase L for select sites in tRNAs and Y-RNAs. To cleave only certain positions and only certain RNAs, RNase L has to detect additional features, such as RNA structures or specific protein-RNA complexes. In the case of RNY4, these mechanisms become more important than the intrinsic preference of RNase L for UN^N sites, promoting cleavage predominantly at the CA^G position.

To understand the molecular origins of RNase L specificity, we tested cleavage of synthetic model RNAs derived from tRNA-His and RNY4. The model substrate derived from tRNA-His contains the complete ASL stabilized by three engineered base pairs. The model RNY4 construct contains the nucleotides surrounding the specific RNase L cleavage site and is predicted to fold into a local stem-loop structure (Fig. 2B). We initially considered this local stem-loop structure a possible reason for the site-specific cleavage of RNY4. Strikingly, neither stem-loop was cleaved at the physiologic site (Fig. 4A). Moreover, the CA^G site in the RNY4 model was completely resistant to RNase L.

**Figure 4.**
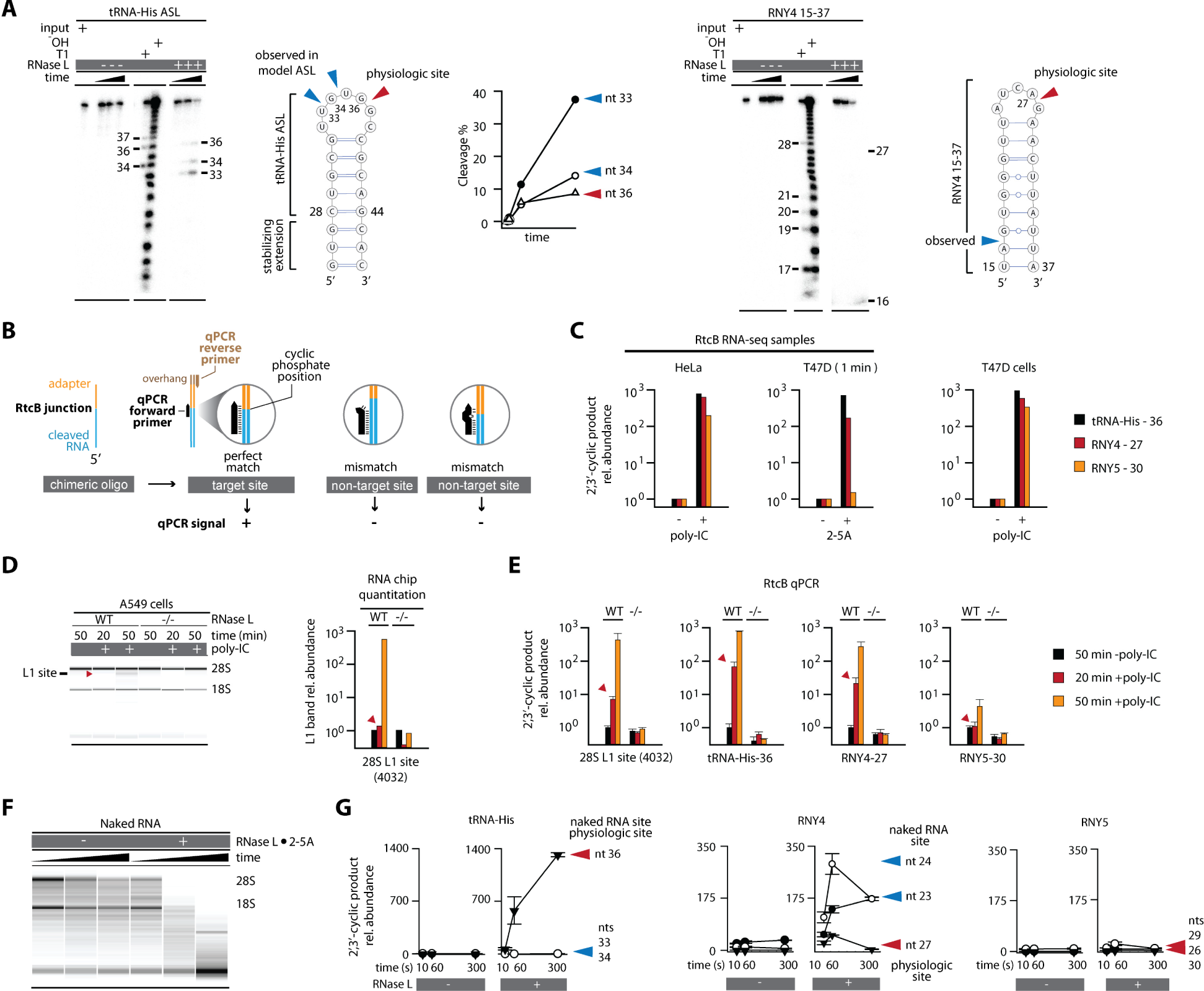
RNase L recognizes structured RNA. (A) Cleavage of model stem-loops from tRNA-His and RNY4 analyzed by polyacrylamide gel electrophoresis (PAGE). (B) A schematic overview of RtcB qPCR. A synthetic adaptor is ligated to RNA cleavage products and the chimeric oligonucleotide is detected by qPCR. (C) RtcB qPCR analysis of RNA samples used for RNA-seq (left panels). Treatment of T47D cells with poly-IC for 8 hours (right panel). (D) BioAnalyzer data from mild poly-IC treatment of WT and RNase L^-/-^ A549 cells. Cleavage of 28S rRNA is not visible at 20 minutes. (E) RtcB qPCR detects robust RNase L activity in the 20-min sample. Error bars show S.E. from two independent RtcB ligations. (F) Cleavage of naked RNA by RNase L. (G) RtcB qPCR analysis of tRNA-His and RNY4 cleavage by RNase L in naked RNA. Error bars show S.E. from two technical replicates.

We next tested whether RNase L can recognize full-length tRNA-His and RNY4. We prepared the full-length RNAs using T7 RNA polymerase and engineered ribozymes to obtain homogeneous oligoribonucleotides, as described previously (Costantino et al., 2008). RNase L cleaved the transcribed full-length tRNA-His selectively in the ASL positions. However, the cleavage targeted all three UN^N sites in the ASL, as seen with the model stem-loop (Fig. S4A; 4A). The cleavage of the transcribed full-length RNY4 occurred predominantly at a non-physiologic site, 24. The physiologic position, 27, was cleaved stronger than many single-stranded UN^N sites nearby, but not as strong as two non-physiologic sites, 23 and 24. Our studies with *in vitro* transcribed substrates thus suggest that specific cleavage of the ncRNAs by RNase L requires cellular machinery.

To test whether host proteins enable the Y-RNA and tRNA recognition, we examined cleavage of naked RNA isolated from human cells. To this end, we developed a highly sensitive qPCR assay for monitoring specific RNase L cleavage in samples of total RNA. This method combines RtcB adaptor ligation with qPCR. The first steps of this RtcB qPCR assay copy the RtcB RNA-seq library preparation. The qPCR step uses a unique site-specific primer that spans the RtcB junction and amplifies the desired cleavage site, but not the neighboring sites (Fig. 4B). RtcB qPCR faithfully detects cleavage of tRNA-His and RNY4/RNY5 in cells (Fig. 4C), and shows a good agreement with the conventional analysis of RNA cleavage by polyacrylamide gel (Fig. S4).

Using this assay we confirmed that the absence of RNY5 cleavage in the semi-permeabilization experiment (Fig. S3C) was not due to an intrinsic inability of T47D cells to cleave RNY5. RtcB qPCR shows a robust RNY5 cleavage in T47D cells in response to poly-IC (Fig. 4C). Next, we compared the sensitivity of RtcB qPCR with the conventional BioAnalyzer readout of RNase L activity based on 28S rRNA cleavage at the L1 stalk (Iordanov et al., 2000; Malathi et al., 2007). To control for RNase L dependence of our measurements, we used WT and RNase L^-/-^ A549 human cells (Li et al., 2016). Transfection of poly-IC for 20 minutes produced no visible rRNA cleavage by BioAnalyzer (Fig. 4D). Nonetheless, RtcB qPCR revealed ~ 10-fold increase in cleavage of the L1 stalk, ~ 30-fold increase in cleavage of RNY4 at site 27, and ~ 100-fold increase in cleavage of tRNA-His at site 36 (Fig. 5E). At 50 minutes, BioAnalyzer and RtcB qPCR both showed a strong cleavage of 28S rRNA, whereas the cleavage of RNY4 and tRNA-His monitored using RtcB qPCR increased to nearly a thousand-fold (Fig. 4E). Our data thus position RtcB qPCR as a next-generation method for analysis of RNase L activation in cells and *in vivo*, which is both sensitive and compatible with complex RNA samples.

**Figure 5.**
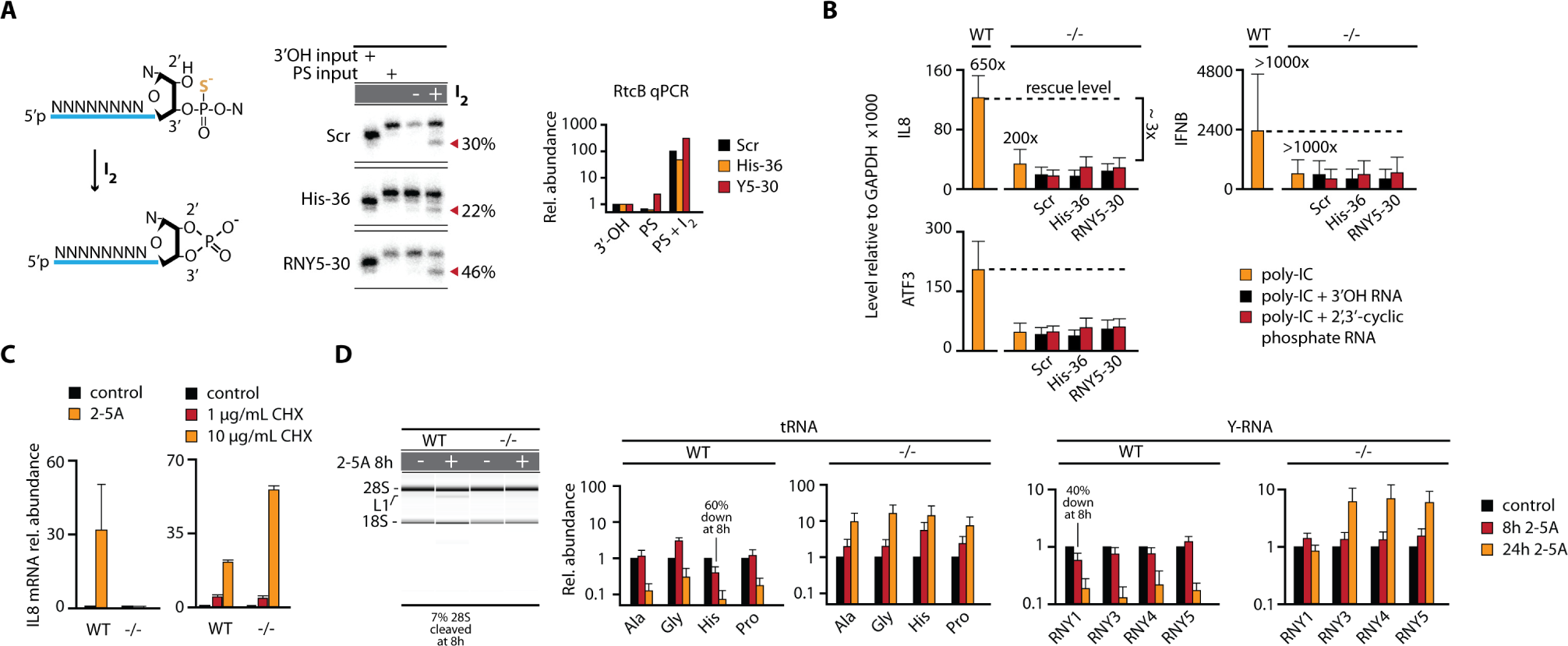
Testing signaling roles of tRNA and Y-RNA fragments. (A) 2’,3’-cyclic phosphate generation in model RNAs with phosphorothioate-iodine. (B) The effect of synthetic 5’-p-tRNA-His and 5’-p-RNY5 fragments with and without 2’,3’-cyclic phosphate on the levels of IFNβ-induced and inflammatory mRNAs. The fragments were transfected for 5 hours into RNase L^-/-^ A549 cells primed with poly-IC for 3 hours.(C) Induction of IL8 mRNA by 2-5A vs cycloheximide treatment of A549 cells for 24 hours. (D) RNase L-dependent downregulation of tRNAs and Y-RNAs by qPCR. Error bars show S.E. from two (cycloheximide, CHX) and three (2-5A) biological replicates.

Here, we used RtcB qPCR to examine whether RNase L can recognize naked tRNA-His and RNY4 in the absence of cellular proteins. Removal of cellular proteins by phenol-extraction changed drastically the overall pattern of RNA cleavage by RNase L. For example, naked 28S rRNA was not cleaved specifically at the L1 stalk and degraded uniformly (Fig. 4F). RtcB qPCR revealed that in striking contrast to *in vitro* transcribed tRNA-His (Fig. S4A), deproteinized tRNA-His isolated from cells is cleaved highly specifically, and at the physiologic position (Fig. 4G). Recognition of tRNA-His by RNase L thus should depend on post-transcriptional tRNA processing by the cellular machinery. The effect of protein removal on the cleavage of RNY4 was markedly different. Deproteinized cellular RNY4 was cleaved non-specifically and resembled the *in vitro* transcribed RNY4 (Fig. 4G; S4A). The critical role of cellular proteins in RNY4 recognition suggests that RNase L targets not the naked Y-RNA, but a protein/Y-RNA particle.

### Probing the Roles of the Y-RFs and tRNA Halves in dsRNA Signaling

RNA cleavage products generated by RNase L were reported to have signaling functions, serving to amplify the IFN response (Malathi et al., 2007) and inflammation (Chakrabarti et al., 2015). Although specific RNAs responsible for this effect have not been identified, it has been proposed that they are small (≤ 200 nt) (Malathi et al., 2007) and contain 2’,3’-cyclic phosphate termini (Chakrabarti et al., 2015). The fragments of tRNA-His and RNY5 identified here represent the most abundant ≤ 200 nt RNA species with 2’,3’-cyclic phosphate released during the dsRNA response, raising the question of their involvement in the IFN signaling or inflammation.

To test this possibility, we obtained a synthetic 5’-phosphorylated oligonucleotide that represents tRNA-His cleaved at position 36 (nucleotides −1 to 36; the nucleotide −1 is the 5’-teminal guanosine added during tRNA-His maturation (Gu et al., 2003)). We also obtained a synthetic mimic of RNY5 cleaved at position 30, which represents an abundant Y-RF (Fig 2B). To create the 2’,3’-cyclic phosphate terminus on a preparative scale, we used a chemical approach based on iodine cleavage of phosphorothioate-substituted RNAs. Here, the starting tRNA/Y-RNA fragments were synthesized with an extra 3’-terminal nucleotide attached via a single phosphorothioate linkage. This linkage was cleaved and converted into a 2’,3’-cyclic phosphate by mild ethanol-iodine treatment (Warnecke et al., 1996) (Fig. 5A).

The tRNA and Y-RNA fragments form naturally during the dsRNA response. Therefore, we examined the effects of the synthetic oligonucleotides in cells primed with poly-IC. We eliminated the background of endogenous tRNA/Y-RNA fragments by using RNase L^-/-^ A549 cells (Li et al., 2016). The combination of RNase L^-/-^ cell background and poly-IC activates all dsRNA surveillance mechanisms, except for the RNase L pathway. If the tRNA/Y-RNA fragments amplify the IFN and inflammatory responses, we expect that i) RNase L^-/-^ cells will produce low levels of IFN-stimulated/inflammatory mRNAs in the presence of poly-IC, compared to WT cells, and ii) supplementation of tRNA/Y-RNA fragments will rescue the deficiency. Poly-IC upregulated IFN-β mRNA normally in WT and RNase L^-/-^ cells (> 1,000-fold), resulting in nearly the same final IFN-β mRNA levels (Fig. 5B). The mRNA of IL8, an inflammatory cytokine involved in NF-κB and NLRP3 inflammasome signaling, was induced normally as well, exhibiting ~ 650-fold increase in WT and ~ 200-fold increase in RNase L^-/-^ cells upon the poly-IC treatment. The levels of a general stress-induced transcription factor, ATF3, were also comparable in poly-IC treated WT and RNase L^-/-^ cells. These results suggest that the RNA fragments released by RNase L contribute no more than ~ 3-fold to the overall ~ 1000-fold IL8 and IFNβ induction by poly-IC. Transfection of the tRNA-His and RNY5 fragments revealed that this contribution is even smaller: the ~ 3.3-fold inflammatory deficiency of RNase L^-/-^ cells was not rescued irrespective of the 2’,3’-cyclic phosphate (Fig. 5B).

Our data suggest that RNA fragments do not amplify the IFN response or inflammatory gene expression, at least in A549 cells. The ability of RNase L to stimulate the expression of IL8 in cells treated with poly-IC was confirmed using direct RNase L activation by 2-5A (Fig. 5C), but this is likely due to translation inhibition because IL8 mRNA is induced similarly by cycloheximide (Fig. 5C). RNase L can inhibit translation by cleaving the ribosomes, as proposed in the early studies (Hovanessian et al., 1979), as well as by cleaving and downregulating tRNAs, which are affected even stronger than the ribosome (Fig. 5D). In summary, the roles of the tRNA and Y-RNA fragments generated by RNase L are presently unclear, although these fragments remain intriguing as candidate cell-specific or paracrine messengers produced in abundance during dsRNA response.

### RNY1 Depletion by RNase L is Sufficient for Initiation of Apoptosis

Concurrently with generating RNA fragments, RNase L downregulates the tRNAs and Y-RNAs (Fig. 5D). The depletion of tRNAs must contribute to the RNase L-mediated inhibition of protein synthesis, whereas the possible effects of Y-RNA depletion are less clear. Y-RNAs entering the blood in complex with the protein Ro60 act as self-antigens and induce autoimmune diseases (Hung et al., 2015). The intracellular functions of Y-RNAs only begin to emerge and point to an essential role of Y-RNAs in cell proliferation (Kowalski and Krude, 2015). Y-RNAs, particularly RNY1 and RNY3, are overexpressed in tumors, whereas knockdowns of these Y-RNAs result in cell growth inhibition (Christov et al., 2008). These findings raise the question of whether Y-RNA depletion is responsible for the anti-proliferative and pro-apoptotic effects (Zhou et al., 1997) of RNase L.

Supporting this model, 2-5A transfection readily kills WT cells and downregulates the Y-RNAs, but has no effect in RNase L^-/-^ cells (Fig. 5D; 6A). However, dying cells activate Y-RNA fragmentation (Rutjes et al., 1999), making it uncertain whether the loss of Y-RNAs in our experiment is the cause or a general marker of cell death. To clarify this connection, we tested whether Y-RNA depletion is sufficient for activation of apoptosis. We downregulated individual Y-RNAs using the antisense DNA/RNase H approach, which allows specific Y-RNA knockdowns (Fig. 6B). Surprisingly, depletion of a specific Y-RNA, RNY1, was sufficient for induction of cell death (Fig. 6C). The knockdown caused cell death in both WT and RNase L^-/-^ cells, indicating that the terminal effect of RNY1 depletion does not require cleavage of additional RNA targets by RNase L. Individual knockdowns of the remaining Y-RNAs had small effects, although RNY5 knockdown was weakly pro-apoptotic.

**Figure 6.**
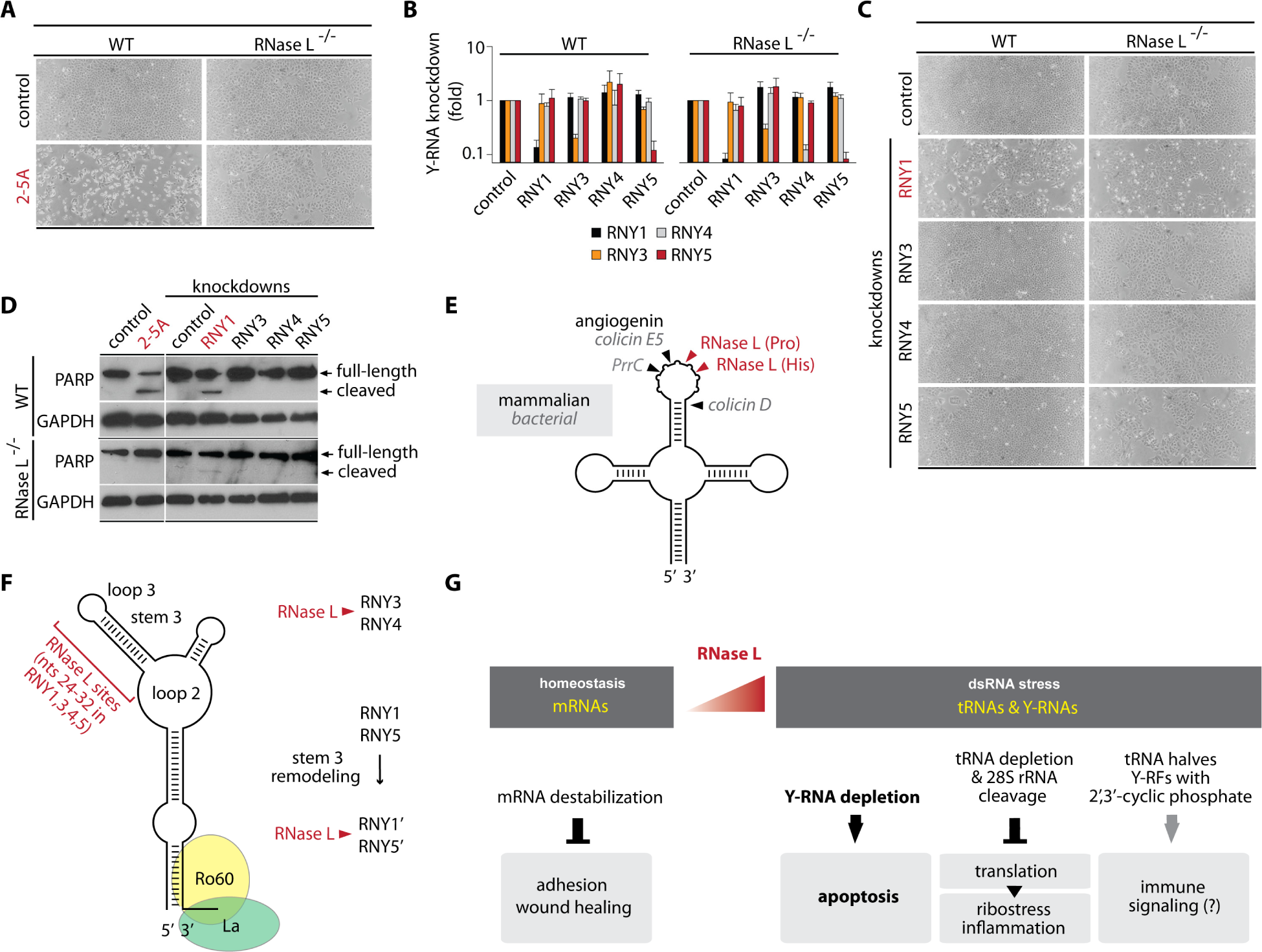
Y-RNA depletion links the dsRNA-RNase L pathway to apoptosis. (A) 2-5A transfection promotes RNase L-dependent death in A549 cells within 24 hours. (B) Knockdowns of individual Y-RNAs by antisense DNA. Error bars show S.E. from three biological replicates. (C) The effect of 24-hour knockdowns of individual Y-RNAs on cell monolayer integrity. (D) Western blot of PARP cleavage indicates activation of apoptosis in dying cells. (E) A graphic summary of tRNA cleavage by bacterial and mammalian endoribonucleases. (F) RNase L cleavage sites in Y-RNAs. Secondary structure of RNY1 is shown for illustration. (G) Suggested mechanism of RNase L signaling in dsRNA response.

To verify that the cells die by apoptosis, we examined the cleavage of poly-ADP ribose polymerase (PARP) (Siddiqui et al., 2015). Consistent with the phenotypes, PARP was cleaved in cells stimulated with 2-5A and in cells with RNY1 knockdown (Fig. 6D). PARP was cleaved in WT and RNase L^-/-^ cells, demonstrating that once RNY1 is downregulated, an RNase L-independent apoptotic cascade is initiated. A stronger PARP cleavage in WT cells suggests that RNase L may amplify the apoptotic response induced by the loss of RNY1. Nonetheless, RNY1 depletion is sufficient to turn on the apoptotic switch without additional RNase L activities.

## Discussion

We have shown that activation of the human innate immune system in the presence of dsRNA promotes cleavage of small ncRNAs by RNase L. By cleaving tRNAs, RNase L functionally resembles the mammalian enzyme angiogenin and the bacterial enzymes colicin E, colicin D, PrrC, zymocin and PaT (Ogawa, 2016). The bacterial enzymes cleave tRNAs to kill competing bacteria and regulate host bacteriostasis. Angiogenin cleaves mammalian tRNAs during cellular stress, such as heat shock or UV exposure, releasing tRNA halves. These tRNA halves have physiologic functions and may inhibit translation initiation, enhance cell proliferation and regulate retroelements (Honda et al., 2015; Ivanov et al., 2014; Sharma et al., 2016). RNase L does not resemble angiogenin or bacterial endoribonucleases structurally and cleaves its tRNA targets at unique positions in the ASL (Fig. 6E). The resulting tRNA halves accumulate near the level of the most abundant cellular RNA with a natural 2’,3’-cyclic phosphate, U6. Whether these tRNA halves have immune or other functions is an intriguing question that awaits future experimental testing.

To the best of our knowledge, RNase L stands out as the only mammalian endoribonuclease reported to cleave Y-RNAs in live cells, raising a possibility that RNase L may have an important role in autoimmune disorders. All major cleavage sites that we detect in Y-RNAs during the dsRNA response belong to a narrow region within the first strand of stem 3 and adjacent nucleotides (Fig. 6F). This regioselectivity suggests that RNase L recognizes a common element found in all Y-RNAs. The analysis of RNY4 cleavage suggests further that RNase L recognizes a protein/Y-RNA complex, which remains to be characterized. The cleavage of RNY3 and RNY4 in stem 3 does not require structural changes because the cleaved bases are in single-stranded regions, whereas cleavage of RNY1 and RNY5 involves remodeling to disrupt stem 3 base pairing. This mechanism controls Y-RNA sensitivity to RNase L (Fig. 2B vs S2B, S3C) and may also regulate the cellular activities of Y-RNAs.

Y-RFs and tRNA fragments are found in abundance in stressed cells, cancer cells and in patient blood (Christov et al., 2008; Dhahbi et al., 2014; Goodarzi et al., 2015; Thompson and Parker, 2009). Our data suggest that RNase L could be responsible for the production of some of these fragments, whereas RtcB qPCR provides a sensitive and specific readout for their detection. The biogenesis of Y-RNA and tRNA fragments by RNase L depends on the presence of intracellular dsRNA, which additionally renders the RtcB qPCR assay a powerful and highly selective sensor for immunogenic dsRNAs in cells and in clinical samples. Whereas detection of IFNs or IFN-induced genes in these samples does not identify the immunogen, detection of the RNase L-derived Y-RNA and tRNA fragments points specifically to the presence of dsRNA.

The mechanism of RNase L signaling that we describe here is specific to the dsRNA response. Under conditions of homeostasis, the action of RNase L is mild and regulates cell adhesion, migration and proliferation by tuning the levels of select mRNAs (Banerjee et al., 2015; Rath et al., 2015) (Fig. 6G). However, the homeostatic functions do not appear to involve tRNA or Y-RNA cleavage because neither is cleaved in unstimulated WT cells stronger than in matching RNase L^-/-^ cells (Fig. 4E: A549 cells; Dataset S1: HAP1 cells). In the presence of dsRNA, RNase L starts cleaving select tRNAs and Y-RNAs and acts as a terminal arm of the innate immune system that eliminates irreparably damaged cells via apoptosis. Multiple RNAs may contribute to the apoptotic cascade, however RNY1 stands out as a mechanistic target of RNase L sufficient for activation of programmed cell death.

## AUTHOR CONTRIBUTIONS

J.D. designed RtcB RNA-seq and RtcB qPCR studies and prepared RtcB RNA-seq reagents and libraries. J.D. conducted biochemical assays. J.D. and S.R. carried out cell biology work and RtcB qPCR analyses. D.K.M generated *in vitro* transcribed ncRNAs. A.K. wrote software for RtcB RNA-seq data processing and analysis. J.D., S.R. and A.K. wrote the manuscript. A.K. supervised the work.

## ACKNOWLEDGEMENTS

We thank Prof. Susan Weiss and Yize (Henry) Li (University of Pennsylvania School of Medicine) for the gift of A546 WT and RNase L^-/-^ cells (Li et al., 2016). We thank Prof. Ileana Cristea Princeton University) for sharing the antibody against human PARP, Prof. Andrei Korostelev for the help in designing ribozyme constructs. We are grateful to Dr. Wei Wang and to the staff at the Princeton University sequencing facility for help with RNA sequencing. We thank members of Korennykh laboratory for reading the manuscript and providing valuable comments.

## Methods

### RtcB RNA-seq Library Preparation

Small miRvana (Life Technologies) purified RNAs from T47D cells (500 ng) or HeLa cells (1 μg) were ligated to 10 μM adaptor, 5’-OH-<u>GAUCGU</u>CGGACTGTAGAACTCTGAAC-3’-desthiobiotin (underlined bases are RNA). The reactions were conducted using 30 μM RtcB, 20 mM HEPES pH 7.5, 110 mM NaCl, 2 mM MnCl_2_, 100 μM GTP, 40U RiboLock RNase inhibitor (Fermentas), 4 mM DTT, and 0.05% Triton X-100 for 1h at 37 °C. The mixture was quenched by 1 volume of stop buffer (8M urea, 1 mM EDTA, 0.1% SDS, 0.02% bromophenol blue, and 0.02% xylene cyanol) and fractioned by 10% PAGE 29:1 with 8 M urea. Gels were stained with SYBR-safe, visualized, and RNA larger than free adaptor was excised from the gel. RNA was eluted from gel slices overnight at 4 °C with gentle mixing in 20 mM HEPES pH 7.5, 100 mM NaCl, 0.05% Triton X-100. Eluted RNA was recovered by ethanol-precipitation with 25 μg glycogen as a carrier.

RNA was reverse-transcribed using MultiScribe Reverse transcriptase (RT) and 2 pmol of primer complimentary to the ligation adaptor. RNA, RT primer, and dNTPs were incubated for 5 min at 65 °C and snap-cooled on ice. A 2X mastermix containing RT buffer, RT, and 40U Ribolock was added to snap-cooled samples for a final volume of 20 μL. Reactions were incubated at 25 °C for 10 min, then at 37 °C for 1h. RNA/cDNA hybrids were pulled down with hydrophilic magnetic streptavidin beads (NEB) and washed three times with 1 mL of 20 mM HEPES pH 7.5, 300 mM NaCl, 0.1% Triton X100, 10 min per wash. This step was followed by two 1 mL washes with 20 mM HEPES pH 7.5, 100 mM NaCl. The final wash buffer was removed and RNA/cDNA hybrids were eluted with 10 μL 10 mM biotin. The eluted cDNAs were then added to an equal volume of 20 mM HEPES pH 7.5, 0.1% Triton X100.

An adaptor, 5’-P-TGGAATTCTCGGGTGCCAAGG-3’-amino, was ligated to the 3’ ends of cDNAs using 1 U/μL CircLigase (Epicentre) and 1 μM adaptor. CircLigase reactions were incubated for 1 hour at 65 °C and quenched by adding EDTA to a final concentration of 8 mM. Quenched CircLigase reactions were PCR amplified for 16 cycles with Phusion DNA polymerase and forward primer with the sequence 5’-AATGATACGGCGACCACCGAGATCTACACGTTCAGAGTTCTACAGTCCGA-3’ plus NEXTFLEX small RNA barcode primers (BIOO Scientific). Libraries were analyzed by Agilent BioAnalyzer high sensitivity DNA 1000 chip. Equimolar amounts were pooled, gel purified, and sequenced on an Illumina HiSeq 2500, such that the sequencing read is the reverse complement of the captured RNA and the first base of the read represents the base that had 2’,3’-cyclic phosphate.

### RNA-seq Computational Analysis

After barcode splitting, sequencing reads with quality scores ≥ 30 were trimmed of adaptor sequences and converted into reverse complements. The obtained reads were mapped to the human transcriptome using FASTA algorithm with ≤ 1 mismatch, which accounts for possible polymorphisms. Sequences of tRNAs were modified to include CCA 3’-ends and G at the -1 position in tRNA-His. All steps past barcode splitting were conducted using software written in-house. Source code and Microsoft Windows binaries are available upon request.

### Tissue Culture

All cell lines were maintained at 37 °C in a 5% CO_2_ humidified atmosphere. HeLa cells were grown in MEM + 10% FBS. A549 WT and RNase L CRISPR KO cells (Li et al., 2016) and T47D cells were grown in RPMI + 10% FBS. HAP1 cells were maintained in IMDM + 10% FBS. HeLa and T47D cells were a gift from the laboratory of Dr. Yibin Kang (Princeton University) and A549 cells were a gift from the laboratory of Dr. Susan Weiss (University of Pennsylvania).

### Cell Treatments for RNA-seq

For poly-IC treatment, HeLa cells at ~ 80-90% confluence were transfected with 1 μg/mL poly-IC or mock transfected (PBS) using Lipofectamine 2000 for 8 hours. Cells were trypsinized, washed with cold PBS and small RNAs were purified using the miRvana kit. For RNase L overexpression, HeLa cells at ~ 80-90% confluence were transfected with 10 μg pcDNA4/TO encoding WT or H672N RNase L (Han et al., 2014) using Lipofectamine 2000. Cells were harvested as above at 24h post-transfection and RNA was purified using the miRvana kit. For 2-5A treatment via semi-permeabilization, T47D cells were trypsinized at ~ 80% confluence and washed with cold PBS. Cells were pelleted by centrifugation at 1000g for 5 min and at 4 °C, and resuspended in 1 mL PBS. Resuspended cells were divided evenly and pelleted again for digitonin semi-permeabilization in the absence or presence of p2-5A_3_ in phosphate buffer, as described (Rath et al., 2015). One half of each sample was added to 300 μL miRvana lysis buffer at 1 min and 3 min time points. Small RNA was purified according to the miRvana protocol.

### Transfections and Cycloheximide Treatment

For all experiments with A549 cells, cells were seeded at a density of 7.5 x 10^4^ cells per well in 12-well plates. Treatments were initiated the following day. All transfections were carried out using Lipofectamine 2000. Cells were transfected with 1 μM ppp2-5A_≥3_ (a mixture of 2-5As with ≥ 3 A monomers) or 1 μg/mL poly-IC for the durations stated in the figures. Synthetic tRNA-His and RNY5 fragments (1 μM) with 3’OH or 2’-3’cyclic phosphate ends were transfected after 3-hour poly-IC (1 μg/mL) transfection. Following poly-IC pretreatment, cells were washed with 2 mL complete growth medium and transfected with synthetic fragments for an additional 5h. Individual Y-RNAs were knocked down by transfecting 100 nM Y-RNA specific or non-targeting anti-sense DNA oligonucleotides for 24h (Table S1). Specific Y-RNA antisense oligonucleotide sequences were previously described (Christov et al., 2006). Cells were treated with 1 and 10 μg/mL cycloheximide or DMSO for 8 hours.

### RNA Purification from Cells

Small RNAs for RtcB RNA sequencing were purified with the miRvana kit (Life Technologies) according to the manufacturer’s instructions. Large RNAs for qPCR were purified using the RNeasy kit (Qiagen), Trizol (Life Technologies), or a modified Trizol protocol in which the aqueous phase was transferred to a clean tube, supplemented with 1/3 volume 100% ethanol and then passed over an RNeasy spin column. RNA bound to the column was processed according to the RNeasy protocol. Small RNAs in the flowthrough were precipitated by adding 1/2 volumes of isopropanol relative to the starting volume of Trizol, washed with 500 μL 75% ethanol, air dried, and resuspended in water. Trace amounts of phenol were removed from the purified small RNAs by sequential extractions with water-saturated 1-butanol and water-saturated diethyl ether.

### Western Blotting

Samples were separated on a 10% BisTris PAGE (NuPAGE), and transferred to PVDF membranes (Life Technologies). Following 15 minute of blocking in 5% milk, the membranes were incubated with 1:2000 rabbit anti-human PARP (Cell Signaling Lot #9542) or 1:5000 mouse anti-human GAPDH (Sigma) primary antibodies at 4 °C overnight. The membranes were then washed with TBST and incubated with horseradish peroxidase-conjugated anti-mouse or anti-rabbit secondary antibodies (1:10,000, Jackson ImmunoResearch) for 30 minutes. The membranes were washed again and detected with ECL Western Blotting Detection Reagents (GE Healthcare Life Sciences) on X-ray film.

### RtcB Enzyme Preparation

RtcB was cloned from *E. coli* into pGEX-6P and expressed with an N-terminal GST-tag. Protein isolation and purification were performed as described for human OAS1 (Donovan et al., 2013), but in the absence of divalent metal cations.

### Synthesis of Model RNAs with 2’,3’-Cyclic Phosphate

Model RNAs were synthesized by Dharmacon and deprotected according to the manufacturer’s instructions. Deprotected RNAs were ethanol-precipitated, resuspended in 20 mM HEPES pH 7.5 and quantified by UV spectrophotometry. Phosphorothioate linkages were converted to 2’,3’-cyclic phosphate in 80-90% formamide, by adding 1/10 volume of I_2_ (1 mg/mL) in ethanol. The reactions were incubated at 37 °C for 10 min. During large-scale preparations, the iodine was fully consumed after 10 min, based on loss of color. Reactions were supplemented with 1/10 volume of 10 mg/mL I_2_ and incubated for another 10 min. There was no loss of color after the second I_2_ treatment. RNAs were ethanol-precipitated, resuspended in 20 mM HEPES, pH 7.5, and quantified by UV spectrophotometry. For analysis by PAGE, small amounts of RNA were dephosphorylated with Antarctic phosphatase (NEB) and 5’-end labeled with PNK and ^32^P-γ-ATP. Model RNA sequences are listed in Table S1.

### Quantitative PCR (qPCR)

Purified small RNAs were reverse transcribed using MultiScribe reverse transcriptase and gene-specific primers (Table S1). qPCR was conducted using SYBR-green based detection over 40–50 cycles. Annealing/extension was conducted at 60 °C for 1 min. U6 RNA was used for normalization. Messenger RNAs were reverse transcribed using the High Capacity RNA-to-cDNA kit (Thermo Scientific). qPCR utilized SYBR-green based detection over 40–50 cycles and annealing/extension temperature of 60 °C for 1 min. GAPDH was used for normalization.

### RtcB qPCR

Small RNAs were ligated to an adaptor with RtcB in 10 μL reactions as described for sequencing library preparation. Reactions were modified to use less Ribolock (10 U). Reactions were stopped by adding EDTA to a final concentration of 3 mM in 11 μL volume and incubated for ~ 10 minutes at room temperature. One μL of EDTA-quenched ligation reaction was used as template for reverse transcription with Multiscribe RT and a primer with a 3’ end that is complimentary to the adaptor and has a 5’-overhang that serves as a universal priming site during qPCR (5’-<u>TCCCTATCAGTGATAGAGAG</u>TTCAGAGTTCTACAGTCCG-3’) (Fig. 4B). Reverse transcription was carried out as described for library preparation, except that 0.4 U Ribolock per reaction and 10 pmol RT primer were used. Reactions were terminated by heating to 95 °C for 5 min. SYBR-green based qPCR was conducted using a universal reverse primer that binds to the cDNA overhang (underlined) and cleavage-site specific forward primers designed for each RNA target. SYBR Green qPCR was carried out for 50 cycles using 62 °C annealing/extension for 1 min. U6, which has a naturally occurring 2’,3’-cyclic phosphate and an RNase L independent cleavage site in tRNA-His (position 18, transcript numbering; Dataset S1) were used for normalization. The *in vitro* digest of naked RNA was normalized using only the internal tRNA-His site 18 because U6 RNA was degraded too rapidly by RNase L. Data for in vitro transcribed RNY4 and tRNA-His cleavage were normalized using the 3’ end of RNY4 and tRNA-His because the HDV ribozyme generates a 2’,3’-cyclic phosphate suitable for RtcB ligation.

### Ribozyme Constructs for RNA Transcription *in vitro*

T7 RNA polymerase transcription constructs of RNY4 and tRNA-His flanked on the 5’ side by the hammerhead ribozyme and on the 3’ side by the HDV ribozyme were synthesized and cloned into pUC57 by GeneWiz (South Plainfield, NJ). Sequences are as follows. The T7 promoter is in bold and the RNY4 and tRNA-His sequences are underlined: RNY4: 5’-GAATTC**TAATACGACTCACTATAGGG**AGAACCATCGGACCAGCCCTGATGAGTCC GTGAGGACGAAACGGTACCCGGTACCGTC<u>GGCTGGTCCGATGGTAGTGGGTTAT CAGAACTTATTAACATTAGTGTCACTAAAGTTGGTATACAACCCCCCACTGCTAAAT TTGACTGGCTTTTT</u>GGGCGGCATGGTCCCAGCCTCCTCGCTGGCGCCGCCTGGG CAACATGCTTCGGCATGGCGAATGGGACCGGATCC-3’; tRNA-His: 5’-GAATTC**TAATACGACTCACTATAGGG**AGATATACGATCACGGCCCTGATGAGTCC GTGAGGACGAAACGGTACCCGGTACCGTC<u>GGCCGTGATCGTATAGTGGTTAGTAC TCTGCGTTGTGGCCGCAGCAACCTCGGTTCGAATCCGAGTCACGGCACCA</u>GGGC GGCATGGTCCCAGCCTCCTCGCTGGCGCCGCCTGGGCAACATGCTTCGGCATGGCGAATGGGACCAATACAATAATAAGCATAATAACCAAGGATCC-3’

### RNA Transcription *in vitro*

Plasmids encoding ribozyme-flanked RNY4 and tRNA-His were linearized with Bam HI (NEB) and transcribed with the MEGAshortscript T7 transcription kit (Thermo Fisher). Transcription reactions were treated with DNase I for 15 min at 37 ºC followed by incubation at 65 ºC for 15 minutes and slow cooling for 20 minutes at room temperature. One volume of 8 M urea gel loading buffer was added to each reaction and samples were briefly denatured at 95 ºC, resolved by 8 M urea 10% PAGE, and visualized by UV-shadowing. Bands were excised, eluted in TE buffer pH 7.5, and precipitated with 3 volumes of 100% ethanol, 1/10 volume 3 M sodium acetate pH 5.2, and 5 μg glycogen. Recovered RNAs were resuspended in H_2_O and quantified by UV spectrophotometry. RNY4 and tRNA-His sizes were verified after 5’-end labeling by 8 M urea 10% PAGE with in vitro transcribed radiolabeled size markers.

### Cleavage of RNA by Recombinant RNase L

RNA from HeLa cytosolic S10 extracts was purified with Trizol. 2-5A and ATP were incubated for 5 minutes at 20 °C in the absence or presence of recombinant human RNase L. Reactions were started by adding RNA (11 μg). Aliquots (5 μL) were quenched by Trizol at 10, 60, and 300 seconds, and RNA was purified. The reactions were conducted in 20 μL volumes using 12 mM sodium phosphate pH 7.0, 140 mM NaCl, 5 mM MgCl_2_, 1.25 mM β-mercaptoethanol, 0.1% Triton X-100, 1 mM ATP, 5 μM p2-5A_3_, and 100 nM RNase L. Synthetic model RNAs were synthesized by Dharmacon and 5’-end labeled with PNK and ^32^P-γ-ATP. The model RNAs were digested in the buffer described above with 100 nM RNase L using 1 μM p2-5A_3_ in 10 μL volume. Aliquots of each reaction were added to formamide gel loading buffer at time points 15, 120, and 600 seconds. Samples were fractionated on 20% 8 M urea PAGE (29:1).

Transcribed RNAs were incubated at 80 ºC for 3 min, supplemented with 10 mM NaCl and 2 mM MgCl_2_, and cooled at room temperature for 20 min. Full-length ^32^P-5’-end-labeled transcribed RNY4 and tRNA-His were digested for 10 minutes and reactions quenched by adding 4 volumes of 8 M urea stop buffer. The RNA samples were analyzed by 15% PAGE with 8 M urea. Unlabeled labeled full-length transcribed RNY4 and tRNA-His (2 μg) were 5’-phosphorylated with cold ATP and PNK, extracted with one volume of acid phenol:chloroform (5:1) (Thermo Fisher), and precipitated with ethanol. Recovered RNA was resuspended in 10 μl H_2_O. RNA (200 ng) was incubated with or without RNase L as above and aliquots at time points 20 s, 3 min, and 10 min were quenched in Trizol. RNA was purified accordingly and resuspended in 10 μl of H_2_O for analysis by RtcB qPCR.

### Error Analysis

For experiments that required repeated measurements, such as qPCR analyses, standard errors (S.E.) were obtained using biological or technical replicates, as stated in figure legends.

